# Deficient thalamo-cortical networks dynamics and sleep homeostatic processes in a redox dysregulation model relevant to schizophrenia

**DOI:** 10.1101/2021.07.20.453026

**Authors:** C Czekus, P Steullet, T Rusterholz, I Bozic, M Bandarabadi, KQ Do, C Gutierrez Herrera

## Abstract

A growing body of evidence implicates thalamo-cortical oscillations with the neuropathophysiology of schizophrenia (SZ) in both mice and humans. Yet, the precise mechanisms underlying sleep perturbations in SZ remain unclear. Here, we characterised the dynamics of thalamo-cortical networks across sleep-wake states in a mouse model carrying a mutation in the enzyme glutathione synthetase gene (Gclm-/-) associated with SZ in humans. We hypothesised that deficits in parvalbumin immunoreactive cells in the thalamic reticular nucleus (TRN) and the anterior cingulate cortex (ACC) - caused by oxidative stress - impact thalamocortical dynamics, thus affecting non-rapid eye movement (NREM) sleep and sleep homeostasis. Using polysomnographic recordings in mice, we showed that KO mice exhibited a fragmented sleep architecture, similar to SZ patients and altered sleep homeostasis responses revealed by an increase in NREM latency and slow wave activities during the recovery period (SR). Although NREM sleep spindle rate during spontaneous sleep was similar in Gclm-/- and Gcml +/+, KO mice lacked a proper homeostatic response during SR. Interestingly, using multisite electrophysiological recordings in freely-moving mice, we found that high order thalamic network dynamics showed increased synchronisation, that was exacerbated during the sleep recovery period subsequent to SD, possibly due to lower bursting activity in TRN-antero dorsal thalamus circuit in KO compared to WT littermates. Collectively, these findings provide a mechanism for SZ associated deficits of thalamo-cortical neuron dynamics and perturbations of sleep architecture.

## Introduction

Schizophrenia (SZ) is a mental illness that diminishes the life quality of the individuals and their relatives on a daily basis. Clinically, the illness is often diagnosed based on the presence of hallucinations, delusions (positive symptoms) or apathy, anhedonia (negative symptoms) in addition to deficits in attention, cognitive abilities, including sensory and emotional processing. Moreover, sleep disturbances are commonly observed in patients, associated with exacerbated psychotic symptoms and poor clinical outcomes ^1-3^. Interestingly, they are often reported during the prodromal phase preceding a first-episode of psychosis ^4,5^. Sleep perturbations include changes in the latency to sleep onset, reduced latency to stage 3 sleep and rapid eye movement (REM) sleep, and fragmentation of the sleep-wake cycle architecture while REM sleep remains unchanged ^6^. Polysomnography studies in SZ patients revealed a non-rapid eye movement (NREM) sleep reduction of slow-waves (SWs) and spindles ^7-10^, in particular in individuals with high-risk of developing psychosis ^1,9,11^.

Abnormal sleep spindles and SWs reflect aberrant coordinated activity within thalamo-cortical networks which has been linked to the neuropathophysiology of SZ ^12,13^. The interplay of thalamic and cortical networks modulates local topography of sleep oscillations including slow wave activity (SWA) and spindles ^14-21^. During NREM sleep, high amplitude SWA nests higher frequency oscillations in humans, cats^22^, and rodents^15^ that result from a precise orchestration of an interplay between inhibitory (GABAergic tone) and excitatory (glutamatergic tone) neurons within thalamo-cortical networks ^23-25^. Accumulation of sleep pressure during wakefulness, or fragmented sleep, dissipate during sleep a phenomenon called sleep homeostasis that is measured by the SWA ^21-23 26,27^. SWAs are more prominent in frontal cortices and it has been linked to cognitive processing including memory consolidation and its correlation with spindles ^1,28-33^, hence, assessing SWA dynamics may be informative of cognitive deficits associated with SZ.

Spindles emerge from thalamo-cortical interactions as transient and distinct brain oscillations (9– 16 Hz) and reflect sleep stability in both rodents and humans ^14,23,24^. In rodents, they are commonly defined as waxing and waning oscillations of variable peak amplitude (∼100 µV) and duration (>400 ms), often coinciding with the UP-state of cortical slow waves or after a K-complex ^34,35^. Spindles are generated by thalamo-cortical interactions between the GABAergic neurons (mostly parvalbumin (PV)) expressing neurons of the thalamic reticular nucleus (TRN) ^20, 36^ and the excitatory cortico-thalamic cells ^31,37,38^. At the cellular level, spindles are generated by the transient burst firing of TRN neurons and thalamo-cortical (TC) relay cells, which generate typical spindling activity within cortico-TC pathways ^10,13,14,16,34, 31,37,38^. Burst firing is triggered by the activation of T-type calcium channels – i.e., Cav3.2 and Cav3.3 in TRN neurons, Cav3.1 in TC cells ^39^ - that synaptically activate layer IV pyramidal neurons in corresponding cortical areas and elicit excitatory postsynaptic potentials and spindle oscillations. Both intrinsic ^13^ and functional inputs to TRN cells from cortical origins are sufficient to generate spindle-like activity ^36^, whereas cortical inputs to the TRN are responsible for spindle termination ^37^. Functionally, a correlative role for spindles in memory consolidation, intelligence, and cognition has been proposed ^25,26,27,28,29^, suggesting that the integrity of spindles is essential to higher brain functions. In fact, temporally extended and compressed spindles are associated with SZ and other brain disorders, respectively ^1,2,12^.

The biological mechanisms underlying sleep disturbances and altered brain oscillations during NREM sleep in SZ remain unclear. Accumulating evidence indicates that redox dysregulation and susceptibility to oxidative stress (OxS) are among the pathological mechanisms driving the emergence of psychosis ^40^. Energy metabolism and redox-dependent processes are intrinsically linked via multiple cross talks to circadian cycle and activity to maintain proper homeostasis ^41-43^. Therefore, we postulated that compromised redox regulation will affect sleep and its homeostatic regulation, as well as TC network dynamics during sleep. To test this hypothesis, we used *Gclm* ^*-/-*^ mice as a model of redox dysregulation relevant to SZ ^44^. These mice have low brain levels of glutathione synthetase (GSH) ^45^, similarly to patients ^46^ and display oxidative stress together with anomalies in PV neurons-associated networks in several brain regions of human subjects, including the TRN and the ACC ^45, 46,47^. In addition, using *ex-vivo* preparations, TRN neurons of *Gclm* ^*-/-*^ mice show impaired bursting density and firing patterns ^46^. Here, we characterized the sleep architecture and the sleep oscillations in *Gcml*^*-/-*^ mice using multisite *in-vivo* electrophysiology in freely-moving mice. We simultaneously recorded cortical, sensory and non-sensory thalamic network dynamics during spontaneous sleep and subsequent sleep recovery. We found that *Gclm* ^*-/-*^ mice showed a fragmented sleep architecture and altered local sleep homeostasis in non-sensory TC networks. Our work provides new brain-wide circuit mechanisms relevant to SZ.

## Methods

### Animals

We used Gclm^*-/-*^ (KO) mice and Gclm^*+/+*^ (WT) littermates from a breeding maintained at the “Centre d’Etude du comportement” at Lausanne University Hospital, originally reported in ^44^. Animals were housed in individual custom-designed polycarbonate cages at constant temperature (22 ± 1 °C), humidity (40-60%) and circadian cycle (12-h light–dark cycle, lights on at 08:00). Food and water were available ad libitum. Animals were treated according to protocols and guidelines approved by the Veterinary office of the Canton of Bern, Switzerland (License number BE 49/17 and BE 18/2020). Criteria for inclusion and exclusion are included in supplemental materials and methods.

### Instrumentation

Animals were instrumented with two EEG electrodes (frontal: AP −2.0 mm, ML +2.0 mm, and parietal: AP −3.0 mm, ML +2.7 mm) together with tetrode wires inserted into the reticular thalamic nucleus TRN (AP −0.8 mm, ML +1.7 mm, DV −3.5 mm), Ventral posterolateral nucleus (VPL) (AP −1.6 mm, ML −1.82 mm, DV −3.6 mm), anterior dorsal thalamus (AD) (AP −0.85 mm, ML +0.75 mm, DV −2.75 mm), sensory cortex (Brr) (AP −1.8 mm, ML +2.8 mm, DV −1.5 mm), anterior cingulate cortex (ACC) (AP +1.2 mm, ML +0.2 mm, DV −1.5 mm). Two bare-ended wires were sutured to the trapezius muscle on each side of the neck to measure electromyography (EMG) signals. Details on the surgical procedures and materials are included in supplemental materials and methods.

### In vivo electrophysiological recordings

For all multisite recordings, mice were connected to a tethered digitizing headstage (RHD2132, Intan Technologies) and data sampled at 20 kHz was recorded with open source software (RHD2000 evaluation software, Intan Technologies). For details see supplemental materials and methods.

### Histological characterization and immunohistochemistry

Confirmation of electrode placement (Figure 2 and suppl. Figure 2) was carried out as described in supplemental materials and methods. Immunolabeling of fast-spiking PV GABAergic interneurons in the ACC and the TRN were labelled as described in supplemental materials and methods and quantified by manually counting the number of PV cells in a selected subarea of the brain structure in a 8 bit image, automatically thresholded and analysed for number of particles using ImageJ software analysis tool.

### Determination of vigilance state

We scored vigilance states manually, blind to the experimental conditions, in 1 s epochs using the concurrent evaluation of EEG and EMG signals and power band analysis. Vigilante states have been defined as previously described and are detailed in supplemental materials and methods. Data analyses were carried out using custom scripts written in MATLAB^®^ (R2018b, MathWorks, Natick, MA, USA). Furthermore, built-in functions from Wavelet and Signal Processing toolboxes of MATLAB were investigated.

### Spectral analysis

Power spectral density (PSD) was estimated using the Welch’s method (pwelch, MATLAB R2018b Signal Processing Toolbox), using 8 s windows having 75 % overlap. A detailed description of the power calculations, spindle detection and quantifications as well as correlation and modulation between frequency bands are described in detail in supplemental material and methods.

### Single unit analysis

Multiunit activity was first extracted from bandpass-filtered recordings (600–4000 Hz, fourth-order elliptic filter, 0.1 dB passband ripple, −40 dB stopband attenuation). Filtering, detection threshold and clustering were performed as previously described in ^24,31^. We visually inspected sorted spikes and excluded clusters with a completely symmetric shape, as noise clusters, or with a mean firing rate < 0.2 Hz from further analysis. Mean firing rate was calculated as total number of action potentials during each condition divided by total time spent in that state and reported as number per second (Hz). Burst firing of single units was detected as a minimum of three consecutive action potentials with inter-spike intervals (ISIs) < 6 ms, and preceded by a quiescent hyperpolarized state of at least 50ms as reported in ^31^.

### Statistical methods

A detailed description of the statistical analyses is provided in supplemental materials and methods.

## Results

### Sleep fragmentation in *Gclm*^*-/-*^ mice

Sleep is commonly affected in human subjects with SZ and appears as prodromal signature prior to the first and recurring psychotic episodes ^10,48,49^. Alterations on sleep onset and sleep maintenance are present regardless patients are treated or untreated. Yet the mechanisms underlying the sleep disturbances are elusive. Here, we compared the sleep architecture of *Gclm*^*+/+*^ *(WT)* and *Gclm*^*-/-*^ *(KO)* mice susceptible to OxS that leads to a deficiency in PV expression, using longitudinal polysomnographic recordings during the dark and light cycle, while baseline conditions and the sleep recovery period after a protocol of sleep deprivation (SD) (see supplemental materials and methods; Figure 1A-E). We used a SD protocol that consisted in keeping mice (both WT and KO) awake for an extended 4-hour period at the onset of the light period and recorded the subsequent recovery sleep (also called sleep rebound, SR) (Figure 1D). We found that KO mice showed an increase in the number of wake and NREM sleep episodes evident in both light/dark cycles during spontaneous sleep-wake cycle, that renormalized after SD (Figure 1F-H). Concomitant to this increase, the wake bout duration was significantly reduced in both phenotypes with less effect on the total amount of vigilant states (suppl. Figure 1A). Interestingly during the first hour of the SR period, we observed a significant increase in the latency to NREM and REM sleep onset in KO mice as compared to WT mice (Figure 1I-J). Together, these data revealed a strong sleep fragmentation in KO mice accompanied with a decrease in NREM sleep consistent with clinical studies in SZ patients ^10,48,49^.

**Figure 1.**
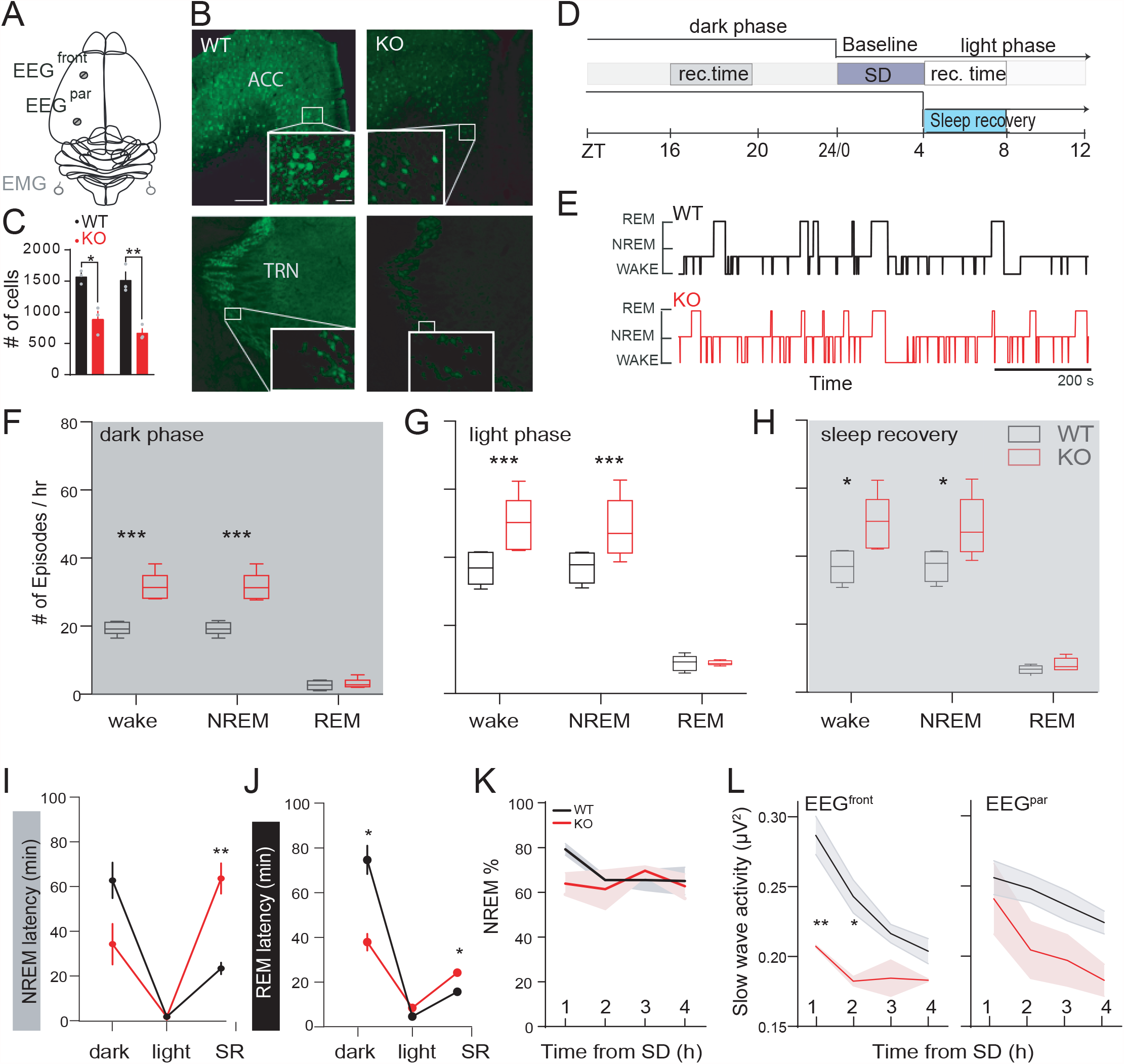
Characterization of Sleep and Sleep homeostatic responses in GCLM^+/+^ (WT) and GCLM^-/-^ (KO) mice. **A**. Schematic representation of electroencephalogram (EEG) electrode placement. **B**. Micrographs showing immune-reactive labelled parvalbumin (PV+) neurons in the ACC and the TRN of adult WT and KO mice. **C**. Significant differences in number of PV+ cells in WT and KO mice. **D**. Experimental recording time line. EEG/EMG signals were recorded for 4 hours in the light period and 4 hours in the dark period to represent wake sleep cycles. A protocol of 4 hours of sleep deprivation (SD) and subsequent SR period were recorded. Data analysed for SR was taken from the first hour of the SR. **E**. Hypnograms taken from a representative wild-type litter mate (WT, black trace) and a GCLM^-/-^knockout animal (KO, red trace). **F**. Min-Max graphs of summary data of the number of episodes per hour +/- s.e.m during the dark n=5,6; for WT and KO respectively; for dark; **P< 0.001, t= 7.3, DF= 27 Sidak’s comparison tested for WT and KO). **G**. For the light period (n=8,7; for WT and KO respectively; **P< 0.001, t= 4.6 for wake and t= 3.98 for NREM sleep, DF=36 for WT and KO). **H**. Data from the first hour of SR (n=7,7; for WT and KO respectively; *P= 0.34, t= 2.66 for wake and *P= 0.14, t= 3.0, DF=36 for NREM tested for WT and KO). **I**. Average latencies to the first consolidated NREM sleep bout +/- s.e.m for WT (black) and KO (red) during the dark, light and SR periods (n=6, 8 and 6 for WT and n= 6,6 and 5 for KO for dark, light and SR phase respectively; * P= 0.007; t= 5.56, DF= 5.16). **J**. Averaged latencies to the first REM sleep episode +/- on the light, dark and SR phases (n=4, 8 and 8 for WT and n= 6,6 and 5 for KO for dark, light and SR phase respectively; *P= 0.027; t= 6.02, DF= 3.0 for dark and *P= 0.036; t= 4.3, DF= 4.0 for SR). **K**. Average NREM sleep total duration +/- s.e.m over 4 hours after extended wakefulness at the onset of the light phase (n= 5 and 5 WT and KO respectively). **L**. Normalized time progression of SWA changes +/- s.e.m on the frontal EEG derivation (left, n= 6 WT and n= 3 KO; **P= 0.009, t= 5.742, DF= 5.049 and *P= 0.039, t= 3.946, DF= 5.282) and from parietal EEG (EEG^par^) electrode showing no significant differences (right). All multiple comparisons were carried out using two-way ANOVA and Sidaks’s multiple comparison test.

Sleep fragmentation is often associated with altered the SWA (dominated by large amplitude and low frequency EEG oscillations) during spontaneous NREM sleep and SR. Interestingly, a time-course analysis revealed no homeostatic changes of SWA during SR in KO as compared to controls (phenotype differences **P = 0.003 F (1, 7) = 19.89). Furthermore, remarkable differences were observed in the topography of these changes (Figure 1L) suggesting a dysregulation of the networks underling the modulation of sleep SWA in KO mice. In order to further investigate this aspect, we sought to characterized the delta oscillations during spontaneous NREM sleep and SR using a multisite electrophysiological approach that allows to study local neuronal activity as well as network interactions. Delta oscillations can be split into δ1 (0.75-1.75), relative insensitive to sleep homeostasis, and δ2 (2.75-3.5) whose amplitude directly depends on the time spent awake and that correlates with activity in the medio-dorsal thalamus and the prefrontal cortex (PFC) ^26^. We recorded cortical EEG signals concurrently with depth local field potentials (LFP) of high order thalamic (ACC), antero-dorsal thalamic nucleus (AD), TRN, ventral posterolateral/medial nuclei (VPL/M) and somato-sensory cortex (Brr) in freely-moving mice (see methods, Figure 2A-B and suppl. Figure 2A). As previously reported ^26^, δ2 amplitude was significantly increased during the SR period in WT mice but not in KO littermates. In addition, delta amplitude in baseline conditions, δ2 was significantly lower in the frontal cortices (in both EEG^front^ and ACC electrodes), AD and to a lesser extend in the somatosensory TC networks in KO as compared to WT (Figure 2D). As expected, δ1 activity during SD was minimally affected in WT and found to be similar in KO mice (suppl. Figure 2B). Note that altered δ oscillations were altered in TRN during baseline sleep (BL; suppl. Figure 2B).

**Figure 2.**
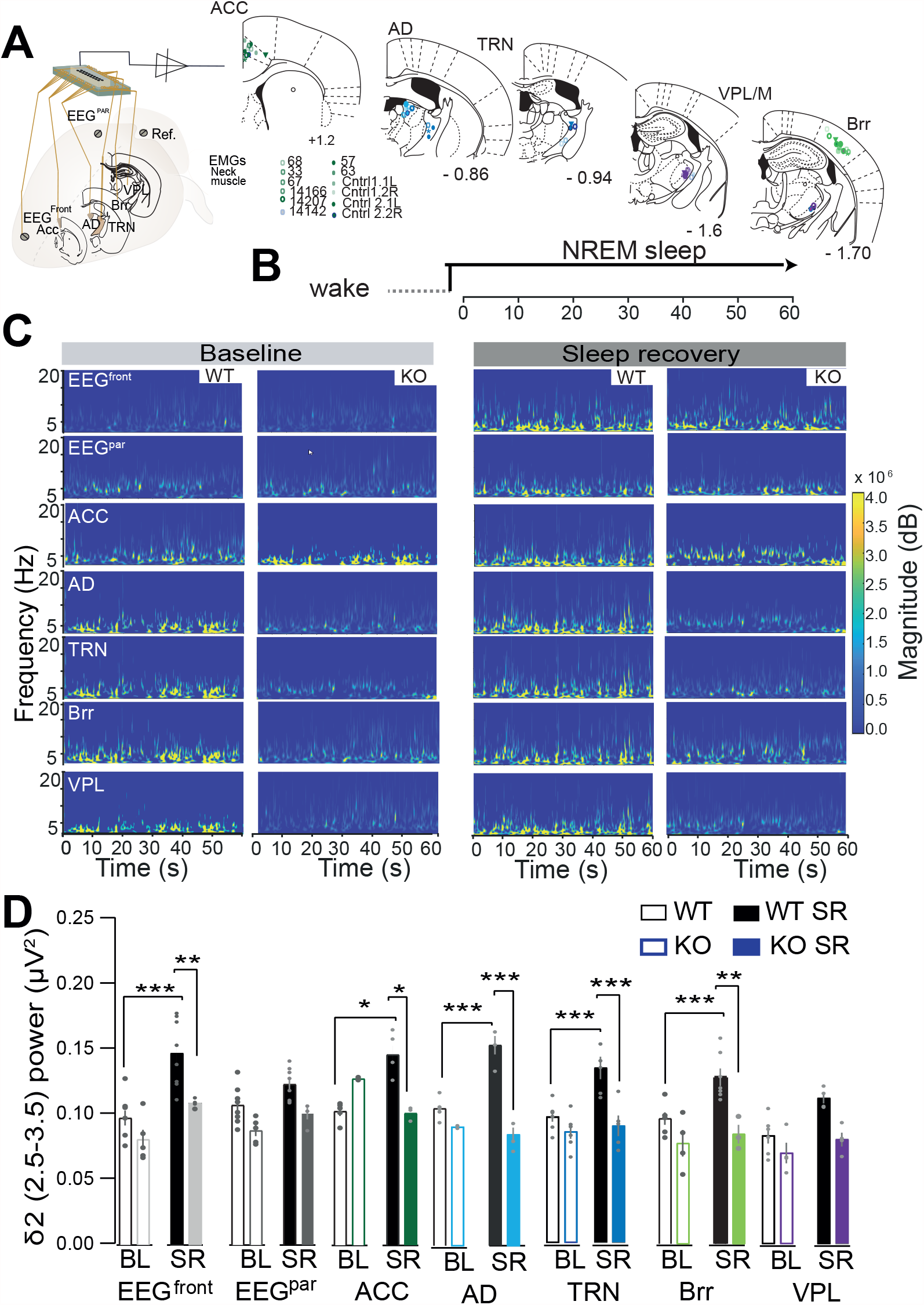
Topographical characteristics analysis of NREM sleep SWA in WT and KO mice in BL and SR period. **A**. *Left*, schematic representation of chronic multisite tetrode implantation for chronic simultaneous EEG, EMG, and tetrode recordings in TC networks: anterior cingulate (ACC), anterior dorsal thalamus (AD), ventral basal thalamus (VPL), reticular thalamic nucleus (TRN) and the somatosensory cortex (Brr) in freely behaving mice. *Right*, schematic of coronal sections taken from *Paxinos and Franklin* ^1^ atlas with electrode locations of all animals included in the analysis. **B**. Experimental timeline for the NREM sleep analysis of the power spectra shown below. **C**. Representative time-frequency of local power spectra recorded from EEG and TC placed electrodes in WT and KO animals during baseline (BL, right) and sleep recovery (SR, left). **D**. Normalized Delta 2 (δ2) power (2.5-3.5) quantification during NREM sleep baseline (BL) and the first hour of sleep recovery time (SR) taken from the EEG and tetrodes in TC nuclei (EEG^front^: WT-WT SR ***P< 0.001, t= 6.67; [WT SR]-[KO SR] **P= 0.002, t= 4.21, n=8 WT and n=5 KO; ACC: WT-WT SR ***P< 0.001, t= 4.12; [WT SR]–[KO SR] **P= 0030, t= 4.62, n=5 WT and n=3 KO; AD: WT-WT SR ***P< 0.001, t= 1.14, [WT SR]–[KO SR] ***P< 0.001, t= 3.01; n=5 WT and n=2 KO; TRN: WT-WT SR ***P< 0.001, t= 4.71; [WT SR]–[KO SR] ***P< 0.001, t= 5.10; n=7 WT and n=5 KO; Brr: WT-WT SR **P= 0.009, t= 3.82; [WT SR]–[KO SR] **P= 0.002, t= 4.25; n=6 WT and n=4 KO; DF= 116, all multiple comparisons were carried out using two-way ANOVA and Bonferroni’s multiple comparison test.

SWs amplitude and slope have been shown to be correlated to the frequency of δ2 in mice ^26^. Using an automated algorithm detection of SWs in mice ^50^, we confirmed that SW slope are increased in cortical and intra-thalamic structures during the first hour of the SR period in WT mice, but absent in KO mice consistent with their impaired sleep homeostasis (suppl. Figure 2C). As mentioned above, redox-dependent processes are linked to circadian cycle and activity to maintain proper homeostasis via multiple cross talks. Thus, we recorded mice during the second portion of the night (ZT 12-16) to test whether changes in the wake inertia or circadian pressure affected delta power. We found no significant changes in the amplitude of δ1, δ2 nor the slope of SWs (suppl. Figure 2D-E) during naturally occurring NREM episodes, suggesting that animals showed normal sleep pressure again once the wake inertia has dissipated ^51^.

### Characterization of spindle dynamics in *KO* mice

To further investigate the influence of OxS (including deficits of PV+ immune positive neurons in the TRN and ACC) of *Gclm*^*-/-*^ *(KO)* mice on TC network dynamics, we quantified the sigma power in both WT and KO mice. Sigma power was not altered in KO mice during the SR period or NREM sleep in BL conditions (suppl. Figure 3A). To eliminate a possible contributions from background EEG activity and selectively quantified spindles event (i.e., within sigma frequency band), we used an automatized detection of sleep spindles ^31^ (figure 3A-B). We observed that spindle rate in spontaneous NREM sleep in WT and KO animals were similar at the EEG levels as well as locally in higher order and sensory thalamic networks (Figure 3C and suppl. Figure 3B, C), however, *KO* mice lacked a homeostatic increase in spindle rate during SD as compared to WT and previously reported ^31^ (WT BL vs KO ***P <0.001, F(3,24)=14.95; KO BL Vs SR P=0.734 (F18,129) = 0.76; Figure 3B-C).

**Figure 3.**
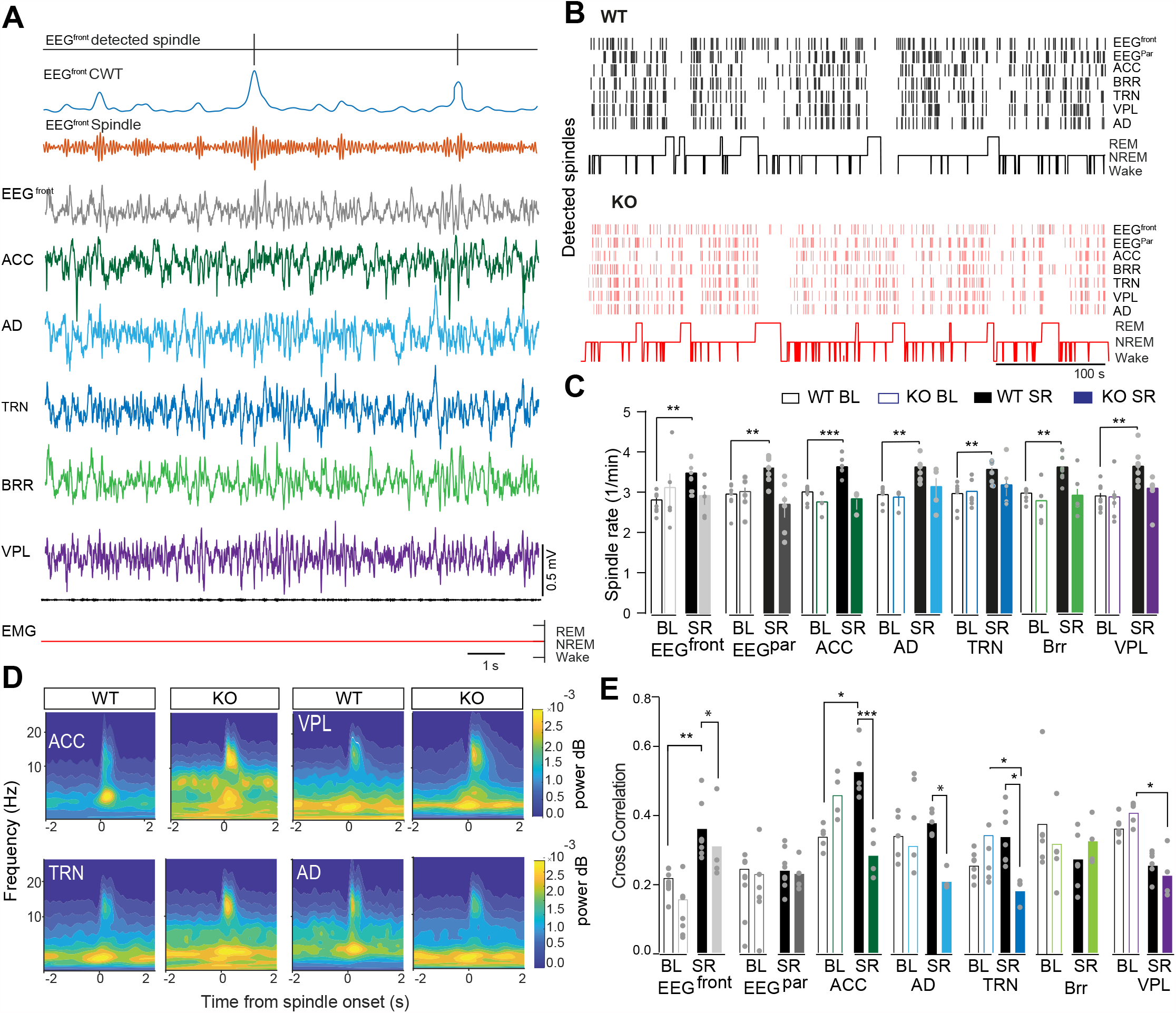
NREM sleep spindles in WT and KO mice. **A**. Representative traces of detected spindles (top), normalized wavelet energy using the complex Morlet and frequency B-Spline functions (blue), filtered EEG signals at spindle range (10-16 Hz, orange) and EEG (grey), LFP traces from ACC (green), AD (light blue), TRN (blue), Brr (light green), VPL (purple) and, EMG (black) and hypnogram on the bottom. **B**. Raster plot of detected spindles in Gclm^+/+^ wildtype (WT) and Gclm^-/-^ knockout (KO) mice through sleep-wake cycles of the light period. Time-frequency analysis of the delta power (0.5-4 Hz) and spindle envelop of the different recorded locations during NREM sleep baseline. **C**. Average spindle density +/- s.e.m during NREM sleep baseline (BL) and sleep recovery period (SR) (significant values from WT-WT SR in EEG^front^: **P< 0.007, t= 4.17, DF= 13.05; ACC: ***P< 0.001,t= 5.79, DF= 13.42; AD: **P= 0.003, t= 4.65, DF= 12; TRN: **P= 0.007, t= 4.11, DF= 13.68; Brr: **P= 0.001, t= 5.6, DF= 10.77; VPL: **P= 0.008, t= 4.04, DF= 13.12; n=8 and 8 for WT and KO respectively, comparisons were calculated using two-way ANOVA with Bonferroni’s multiple comparisons test). **D**. Averaged cross-correlograms of time–frequency representations of spindles and SWA aligned to the peak of the spindle envelop. **E**. Normalized averaged cross correlation during baseline (BL) and sleep recovery (SR) in all recorded locations. (EEG ^front^: WT-WT SR *P= 0.015, t= 3.078, n=8, 7 for WT and KO; ACC: WT-WT SR *P= 0.010, t= 3.204; [WT SR]-[KO SR] ***P< 0.001, t= 3.89, n=5 WT and n=3 KO n=5,4 for WT and KO; AD: [WT SR]-[KO SR] *P= 0.032, t= 2.83, n=5 and 3 for WT and KO; TRN: KO-KO SR *P= 0.36, t= 2.80, n=7 and 7 for WT and KO; VPL: KO-KO SR *P=0.024, t=0.48, DF= 129, two-way ANOVA with Bonferroni’s multiple comparisons test).

Differences in SWA amplitude and spindle dynamics *Gclm*^*-/-*^ *(KO)* mice are suggestive of possible alterations in the interactions between oscillations in different frequency bands. To assess the relation between SWA and spindles, we extracted SWA (0.5–4 Hz) and spindle envelopes from all signals recorded using multisite LFP recordings (Figure 3D) and computed the SW–spindle CFC using the normalized cross-correlation analysis between the peak of the SW and spindle envelopes for all spindles occurring during NREM sleep (see supplemental materials and methods). Consistent with previous reports ^31^, we observed an increase in CFC in ACC after sleep deprivation in WT (ACC: P= 0.034; q = 5.36; DF = 6.02; two-way ANOVA with Tukey’s post-hoc test) whereas KO mice showed the opposite dynamics (KO ACC * P= 0.03, t = 5.30, DF = 7.22; Figure 3E). Note that CFC during SR is significantly different between phenotypes as well in AD and TRN, suggesting that the frontal and higher order thalamic but not somato-sensory thalamic networks were affected in *Gclm*^*-/-*^ *(KO)* mice (Figure 3E).

We next quantified the phase amplitude coupling (PAC) reflects oscillation synchrony associated with cognitive and sensory processing in humans ^52,53^ and rodents ^54-56^ possibly associated with altered procedural and emotional processing in SZ patients ^57-59^. Here, we assigned directionality on the modulation of two different brain areas by determining PAC between cortico-cortical (ACC-BRR), higher order (ACC-AD), intra-thalamic (TRN-AD) and sensory TC (VPL-Brr) pairs (Figure 4A). During spontaneous sleep, we observed a remarkable increase in PAC between ACC-AD and TRN-AD in KO in KO as compared to WT mice (ACC-AD: *P=0.023, t= 2.9, df=86; TRN-AD: **P = 0.009, t = 5.42, df = 6.254; two-way ANOVA with Bonferroni’s multiple comparisons test) but not in other pairs (Figure 4A and C). During SR, a significant increase was observed in ACC-Brr and ACC-AD in KO in comparison to WT mice (ACC-Brr: ***P < 0.001; t = 5.66; ACC-AD: ***P< 0.001, t= 6.06; DF = 86; two-way ANOVA with Bonferroni’s multiple comparisons test, figure 4B and C). These data further confirm the specific alterations of higher order TC network dynamics in KO mice and suggest an increase in synchronization of non-sensory TC networks.

**Figure 4.**
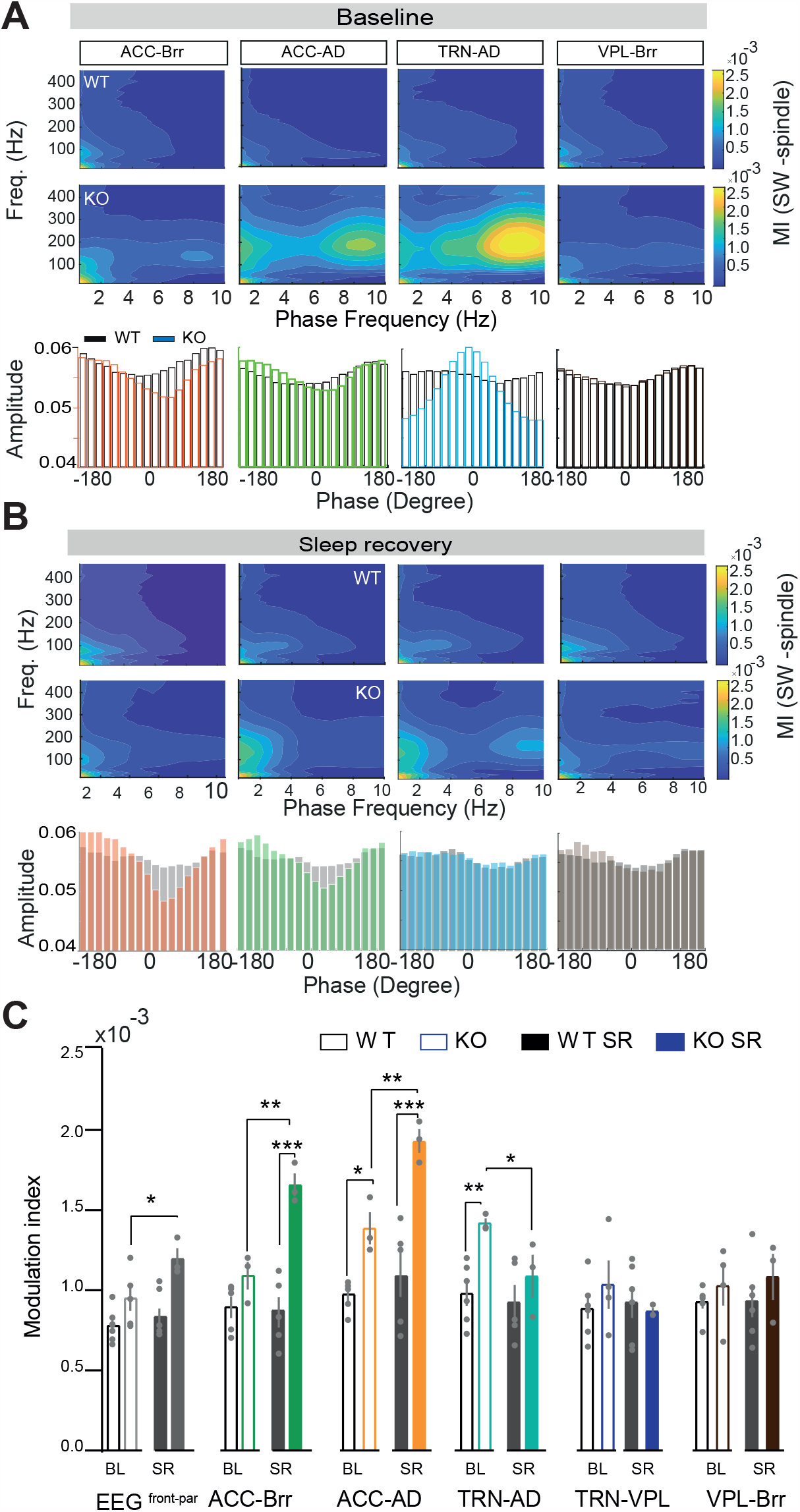
Topographic analysis of brain dynamics in Gclm^-/-^ KO mice. **A**. Averaged phase-amplitude comodulograms (0.5-4 Hz phase frequency and spindle range 10-16 Hz amplitude frequency) computed for connected TC areas: ACC-AD, ACC-Brr, TRN-AD, VPL-Brr local field potentials (LFPs) during NREM sleep in the light phase (BL) and the first hour of sleep recovery period (SR) (n=8 WT and n=7 KO). Bottom: Normalized phase degree histograms of WT (grey) and KO (coloured). **B**. Average modulation index +/- s.e.m. during NREM sleep in the light phase computed for EEG front and EEG parietal signals, ACC-AD, ACC-Brr, TRN-AD, VPL-Brr TRN-AD, TRN-VB, VB-Brr, ACC-Brr and ACC-AD LFPs (WT n= 6,7,6,5 and 5 and KO n= 3,4,4,3 and 3 respectively. TRN-AD: **P= 0.007, t= 5.421, DF= 6.254). **C**. Average modulation index +/- s.e.m during NREM sleep (BL) and the first hour of the sleep recovery period (SR) for ACC-Brr, ACC-AD, TRN-AD, TRN-VPL and VPL-Brr LFPs (WT n=5,6,6,5 and 5 and KO n=3 respectively. EEG front-par: *P= 0.04, t= 2.78; ACC-Brr: [KO BL]-[KO SR] **P= 0.003, t= 3.67; [WT SR]-[KO SR] ***P< 0.001, t= 6.1; ACC-Brr: [WT BL]-[KO BL] *P= 0.02, t= 2.98; [WT SR]-[KO SR] ***P< 0.001, t= 6.1; [KO BL]-[KO SR] **P= 0.004, t= 2.78; TRN-AC: [WT BL]-[KO BL] **P=0.009, t= 3.3, [KO BL]-[KO SR] *P= 0.04, t= 2.8; DF= 86,significance levels were calculated using 2-way ANOVA and Bonferroni’s multiple comparisons test. All results are represented in mean +/- s.e.m. *P< 0.33, **P< 0.002 and ***P< 0.001).

### Altered spike dynamics in KO mice in non-sensory thalamic circuits

Finally we aimed at elucidating the underlying cellular mechanisms associated with the alterations reported above in KO mice as suggested by reduced propensity to generate bursts of action potentials and decreased T-type calcium currents observed *ex vivo* ^46^. We characterized the spiking activity of thalamic and cortical neurons across vigilant states in WT and KO mice (Figure 5). Examination of firing rates from isolated neurons recorded from WT and KO mice revealed that spiking activity was lower for TRN neurons during transitions from wakefulness to NREM sleep (Figure 5A and B and suppl. Table 1) and higher in the ACC and AD during the same transitions (suppl. Figure 4A) in KO mice as compared to WT animals (TRN: 25.1 +/-2.0, 15.5 =/- 2.9; ACC: 9.08 +/- 1.9, 23.34 +/- 3.8; AD: 4.63 +/- 1.36, 10.26 +/-1.39 for WT and KO respectively). Differences in spiking activity were most prevalent during NREM sleep. Indeed, neuronal spiking rate activity was lower in TRN and in ACC in KO as compared to WT animals (TRN: 24.2 +/- 4.05 and 8.45 +/- 1.45; and ACC: 5.36+/- 0.45 and 13.24 +/- 1.92 WT and KO respectively) but showed a renormalisation during SR. Interestingly, this was possibly due to a lower proportion of bursting cells (i.e., less than one burst/min, see methods; Figure 5G). Consistent with our results above, neuronal burst densities were decreased in TRN and Brr, but increased in ACC, when comparing KO to the WT animals (TRN: 31.27 +/- 4.73 and 9.73 +/- 2.22; Brr :26.05 +/- 6.34 and 1.97+/- 0.31; and ACC: 1.66 +/- 0.19 and 4.98 +/- 0.42 # of bursts/min for WT and KO respectively; Figure 5H), suggesting a change in the excitation-inhibition dynamics in both TC loops. The burst density has been previously shown to be affected in KO mice in *ex-vivo* preparations ^46^ and reported to be modulated by the balance between inhibitory and excitatory interplay within the TC networks while he length of a burst is determined by cortical influences ^60^. In fact, we found that burst density amongst TRN -AD, but not TRN – VPL, circuits were significantly lower in KO as compared to WT animals, consistent with impaired homeostatic regulation of TC networks, in particular amongst AD circuits (8.81 +/- 0.22 BL, 7.28 +/-0.31 SR in ms; Figure 5I). Altogether, these findings suggested a dysregulation of local cellular activity in higher order thalamic networks responsible for the disturbances of fronto-parietal oscillatory activity during sleep.

**Figure 5.**
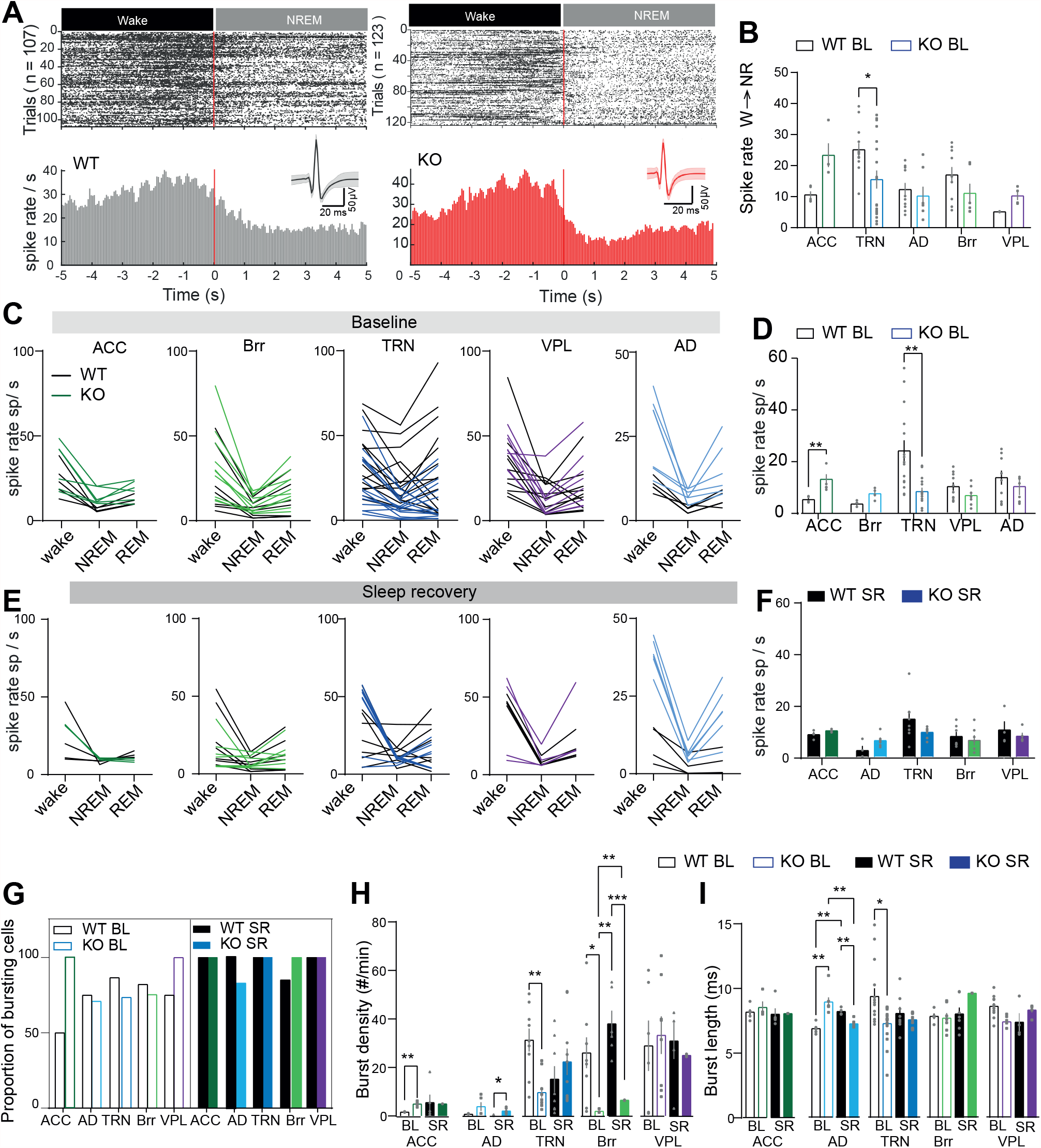
Changes in spiking activity in Gclm^-/-^ KO mice. **A**. Raster plot spiking activity during wake to NREM transitions isolated from tetrode implanted in the TRN in WT (107 transitions) and KO (123 transitions). Bottom, histogram of spiking rate per sec during the transition and representative isolated unit from WT (black) and KO mouse (red). **B**. Summary data +/- s.e.m of spiking rate in transitions from wake to NREM sleep in ACC for WT and KO (n= 5, 4), Brr (n=10,7), TRN (n=13,19), VPL (n=10,6) and AD (n=2,4), significance level * P= 0.018, t= 3.03, DF= 70). **C**. Average spiking rate +/- s.e.m in wake NREM and REM in in all recording sites of all isolated spiking neurons. **D**. Summary data of spiking rated in NREM sleep during the baseline recoding ACC for WT and KO (*P= 0.036, t= 3.64, DF= 17.86, n=5,6), Brr (n=10,7), TRN (n=15,14 **P= 0.009, t= 3.9, DF= 5.61), VPL (n=10,10) and AD (n=5,4). **E**. Spiking rate responses after SD during the first hour of sleep recovery (SR) and summary data (**F)**, same n values that in E. **G**. Proportion of all recorded cells in the different TC nuclei where bursts were detected during the NREM sleep BL and during the SR. **H**. Averaged burst density during NREM sleep baseline and SR recorded(ACC: WT-KO **P= 0.005, t= 7.18, DF= 4.98, n=2,5 for WT and KO; TRN: WT-KO **P= 0.009, t=4.12, DF= 11.43, n=9,10 for WT and KO; Brr: WT-KO **P= 0.031, t= 3.8, DF= 8.04; [KO SR]-[WT SR] **P= 0.007, t= 6.66, DF=5.03; [KO BL]-[KO SR] ***P< 0.001, t= 14.14, DF. 5.75; [WT SR]-[KO SR] *P= 0.013, t= 5.828, DF= 5.003) **I**. Averaged burst length during NREM sleep baseline and SR recorded (AD: WT-KO ***P< 0.001, t= 14.08, DF= 6.32, [WT BL]-[WT SR] **P= 0.005, t= 8.4, DF= 5.88, [KO BL]–[KO SR] ***P< 0.001, t= 9.87, DF= 9.01; [WT SR]-[KO SR] *P= 0.34, t= 5.18, DF= 7.3; TRN: WT-KO *P= 0.046, t= 3.84, DF= 15.30. ACC [BL n=2,5, SR n=5,3], Brr [BL n=6,9, SR n= 6,4], TRN [BL n=9,10, SR n=8,10], VPL [BL n=6,8, SR n=5,5] and AD [BL n=4,5, SR n=2,6]. Significance levels were calculated using two-way ANOVA and Bonferroni’s multiple comparison test. Data is represented in +/- s.e.m. Significance levels represent *P< 0.33, **P< 0.002 and ***P< 0.001.

## Discussion

In this study we investigated for the first time the TC modulation of local and brain-wide dynamics in the *Gclm*^*-/-*^ mouse model of SZ using *in-vivo* electrophysiology in freely-behaving mice. First, we found that *Gclm*^*-/*-^ mice showed deficits in PV+ immunnoreactive cells in the TRN and ACC as previously shown ^46^ and a significant fragmentation on the sleep-wake cycle due to an increase in the number of sleep and wake episodes, both in the inactive (light) and the active (dark) period and SR. Features of sleep homeostasis, including latency to first NREM bout, delta power and spindle density were impaired in KO animals during SR period (Figure 1). These results are consistent with sleep disorders present in some SZ patients ^1,9,49^ as well as the increased sensitivity to mild stress situations reported in this mouse model ^61^.

While circadian influences modulate characteristics of SWs ^63^, sleep-dependent modulation of SWs during NREM is prominent in frontal cortices and implicate non-sensory TC circuits including the medio-dorsal thalamus in mice ^26,64-66^ and humans ^10,67^. Although the TRN neurons have been implicated in spindle generation in somato-sensory thalamo-cortical areas ^17,31,68^, their synaptic contact onto medio-dorsal thalamus and its targets in the ACC is hypothesized to influence the synchronisation of SWs in frontal area in a brain wide manner ^15,31^. Furthermore, the propagation of frontally-generated SWs throughout the brain in humans ^25^ and mice require a functional higher-order anterior thalamic nucleus (e.g., AD) ^24^. Interestingly, this fronto-parietal network dynamics showed aberrant activity in subpopulations of SZ patients as revealed by functional connectivity MRI ^69^, including the prefrontalcortex (PFC), anterior cingulate cortex (ACC) and thalamus ^69^. Consistent with this finding, delta activity and single SW analysis in our study showed a fronto-parietal disturbance in the KO suggestive of a dysregulation of the higher order thalamo-cortical networks. This is further supported by the changes in δ2 (2.5-3.5 Hz), which implicate the centro-medial thalamic network (CMT-ACC-AD) ^26^. In fact, KO mice showed a remarkable lack of proper homeostatic regulation of sleep oscillations responses in frontal cortical (ACC and AD) and local thalamic circuits (TRN) dynamics during NREM sleep, where neuronal networks are essential for sleep homeostasis and modulation of SWA ^16,36,64,70^. To note, others TC circuits (VPL, Brr and ACC) also show altered response to the SD, confirming a dysregulation of intra-thalamic and cortico-thalamic networks. Differences in the modulation of δ1and δ2 in the different thalamic and cortical nuclei of *Gclm*^*-/-*^ *(KO)* further supports the idea that δ1 and delta δ2 are modulated by, and recruit, different TC networks. Here, due to low PV+ immunoreactive cells within the TRN and the ACC, it is possible that local changes in the neuronal excitability, as shown in ^71^ may play a role in the input-output ratios at each topographic location, and thus, in the orchestration of TC networks, although this remains to be further investigated.

In this respect, SWs are temporally linked to the expression and topography of spindle occurrence in humans and rodents ^24,25,66^. Coherence between SWs and spindles are affected in the frontal cortex of the KO mouse model, consistent with a role of CMT-ACC-AD circuit in SW propagation in rodents ^24^, and the reduced volume of the medial dorsal (MS) thalamus and MD-PFC connectivity in resting state fMRI from chronic and early SZ patients ^3^. Altogether these findings evidence a pathological lack of temporal coordination of SWs and spindles during NREM sleep in SZ due to reduced MD-PFC connectivity.

Interestingly, recent investigations have pointed out deficits in the TC networks as an underlying mechanism for the deficit in sleep spindles in patients with SZ. Here, despite reduced PV+ immunoreactive neurons and weakened perineuronal networks in the TRN together and its altered intrinsic firing associated with reduced function of T-type calcium channels in KO mice ^46^ (and accompanying manuscript), spindle activity in this mouse model remains similar to control littermates. Yet, KO mice showed a lack of increase in cortical and local TC spindle rates during the SR possibly due to altered modulation of TRN inputs-outputs circuits ^14,16^, in particular the TRN-AD circuit implicated in spindles generation and NREM-to- REM transitions ^31^, although causal implication remains to be investigated. Nevertheless, the absence of spindle alterations during spontaneous sleep may be due to compensatory network dynamics during development in response to altered function and integrity of the PV cells in the KO model ^20^. Furthermore, spindle deficits may not be simply and only modulated by the activity of the TRN cells, but implicate complex network interactions and plasticity processes during NREM sleep that directly depends on the inhibitory or excitatory tone amongst TC feedforward circuits and cortico-thalamic feedback loop’s interplay as previously suggested ^72^. In fact, data collected from animals carrying a mutation in *GRIA1* gene, another mouse model relevant for neuropsychiatric disorders, showed altered glutamatergic transmission and reduced synaptic plasticity, together with deficit in spindle density ^73^. Thus, further investigations of the developmental, plastic and functional alterations associated with the different cell-types of specific thalamic networks are required to better understand the pathophysiological mechanisms involved.

Nonetheless, our data provide with important links between the redox oxidative stress dysfunction and homeostatic changes in SWA and spindles. Underlying mechanisms may include network and cellular activity for the synchronization of brain activity during homeostatic changes including sleep homeostasis. Here, we found that TRN spiking/bursting activity is decreased in the transitions from wakefulness to NREM sleep which, together with the increase in the spiking/bursting activity in the ACC-AD during Wake to NREM transitions, may result in the sleep fragmentation observed in KO mice. One possible mechanism underlying this sleep instability may be related to the activity of T-type calcium channel, as previously suggested by ^46,74,75^. Together with our findings, these suggest that alteration in the T-type calcium currents may alter network dynamics that are reflected at the functional level during sleep modulation, and that may be play a pivotal role in the sleep modulation and homeostasis.

Non-sensory and higher-order thalamic nuclei, such as the medio-dorsal, centro-lateral, and sensory-motor (posterior group and latero-dorsal) nuclei, have distinct patterns of connectivity with prelimbic cortex and other subcortical structures ^1,12,39^. To test whether these could be involved in the pathophysiological mechanisms of *Gclm* ^*-/-*^ *KO* mice, we characterized the phase frequency coupling of delta-spindle oscillations given the extensive evidence suggesting that perceptual grouping, attention dependent stimulus selection, working memory, and consciousness are associated with synchronized oscillatory activity in the theta-gamma but also the SWA-spindle band ^76^. Our results demonstrate that KO mice lack of proper synchronization of sleep oscillatory activity, which may be contributing to some of the behavioural traits reported for these mice ^61^. Yet, our findings warrant further investigation linking neuronal network activity, sleep oscillations and behavioural phenotyping to better understand brain dynamics during different cognitive tasks relevant to SZ.

Growing body of literature highlights the potential of sleep as a window for neuromodulation therapy in neurological and neuropsychiatry disorders. Identification of temporal and topographical alterations in neuronal activity are thus essential to select proper targets for neuromodulation. Recent investigations both in humans ^77,78^ and animal models ^79,80^ have suggested perturbational procedures to enable real-time tracking and manipulation of sleep EEG oscillatory activity using close-loop transcranial stimulation or auditory stimuli to enhance or diminish SWs and spindles and facilitate brain plasticity and repair ^50,78,81^. Our work identified the AD thalamus, as one of the synaptic targets of the TRN, connected to the ACC, that may represent a target for future neuromodulation approaches. Ultimately this will open new ways to identify sleep oscillations as predictive, diagnostic, or prognostic biomarkers for SZ and other psychiatric disorders.

## AUTHOR CONTRIBUTIONS

Author contributions: C.G.H. and D.K. Q. conception and design of research; C.G.H performed experiments; T.R. and I.B. wrote and adapted Matlab custom scripts for data analysis, M.B. wrote spindle detection algorithm, C.G.H. and C.K. analysed data; C.G.H. interpreted results of experiments and prepared figures; C.G.H drafted manuscript; C.G.H, P.S, D.K.Q. and C.K edited and revised manuscript; C.G.H. approved final version of manuscript.

## Acknowledgements

We thank the Centrum for Experimental Neurology for hosting our research and providing with technical support. To Prof Widmer for the use of the fluorescence microscope. This work a were supported by University of Bern Interfaculty Research Cooperation “Decoding Sleep” and SNF 320030_179565 / 1 (Gutierrez Herrera, C.).

## Data availability

All data and scripts are available upon request.

## Footnotes

1 G Paxinos and KBJ Franklin, *Paxinos and Franklin’s the Mouse Brain in Stereotaxic Coordinates*, 2019.

