## Supplemental Materials and Figures for "Deficient thalamo-cortical networks dynamics and sleep homeostatic processes in a redox dysregulation model relevant to schizophrenia"

***Supplemental Material and Methods***

*Animals*

Only adults (>10 weeks old) male mice were used in the experiments. Animals were housed in IVC cages in groups of 2–5 before instrumentation. After implantation, all mice were housed individually. Animals were habituated to the recording cables in their open-top home cages (300 × 170 mm) and kept tethered for the duration of the experiments. Animals were allowed to move freely in the cages during *in-vivo* electrophysiology experiments. Before commencing experimental recordings, baseline sleep was recorded and compared to previously published results [9,38]. Experiments were performed during the “light phase”, the “sleep recovery period” (12:00 (ZGT4)–16:00 (ZGT9)) and the ”dark phase” (24:00 (ZGT16)-04:00 (ZGT20)). All recordings were performed from 16-20 weeks of age. Animals with heterozygote genetic background were excluded.

*Instrumentation*

Animals were anaesthetized with a multimodal balanced protocol consisting in an injection of fentanyl (0.05 mg/kg; Fentanyl, Kantonsapotheke Zurich, CH), midazolam (5 mg/kg; Dormicum®, Roche Pharma AG, Reinach, CH) and medetomidine (0.5 mg/kg; Dorbene®, Graeub AG, Bern, CH) mixed in NaCl (0.9%) to a volume of 3 µl/g body weight. Intraoperative analgesia was provided by a local anaesthesia of lidocaine (Xylocaine 2%, diluted 1:1 in sterile NaCl 0.2 mL/site) on the incision site. Saline 10 ml/kg and meloxicam 5 mg/kg were given subcutaneously. The skin on the head was shaved and aseptically prepared. A single longitudinal midline incision was made from the level of the lateral canthus of the eyes to the lambda skull suture. Two stainless steel screws were placed in the skull to measure EEG (frontal EEG: AP −2.0 mm, ML +2.0 mm, parietal EEG: AP −3.0 mm, ML +2.7 mm) and a third stainless steel screw was place in the cerebellum as a reference. Two bare-ended wires were sutured to the trapezius muscle on each side of the neck to record EMG. Tetrodes were made from four strands of 10 µm twisted tungsten wires, connected to an electrode interface board by gold pins and inserted into the reticular thalamic nucleus (TRN:AP −0.8 mm, ML +1.7 mm, DV −3.5 mm), Ventral posterolateral nucleus (VPL: AP −1.6 mm, ML −1.82 mm, DV −3.6 mm), anterior dorsal thalamus (AD: AP −0.86 mm, ML +0.75 mm, DV −2.75 mm), sensory cortex (Brr: AP −1.7 mm, ML +2.8 mm, DV −1.0 mm), anterior cingulate cortex (ACC: AP +1.2 mm, ML +0.2 mm, DV −1.5 mm,).They were secured to the skull with dental acrylic (C&B Meta–bond). Finally, the implant was stabilized using a methyl methacrylate cement and the animals were allowed a minimum of 5 days to recover in the home cages on top of a heating mat before starting recordings.

*In-vivo electrophysiological recordings*

For all electroencephalogram (EEG), electromyograms (EMG) and tetrode recordings, mice were connected to a tethered digitizing headstage (RHD2132, Intan Technologies) and data sampled at 20 kHz was recorded in an open source software (RHD2000 evaluation software, Intan Technologies). Habituation to the cables was performed progressively from 1 hour the first day, 4 hours the second day, 8 hours the third day and kept plugged for the rest of the recording period. Baseline recordings started 3 days after animals had nested and resumed a normal sleep–wake cycle. All baseline and recovery sleep recordings were performed between 12:00 (ZGT4) to 16:00 (ZGT8). For sleep deprivation, gentle handling was performed when animals were stationary to prevent sleep between 8:00 (ZGT0) and 12:00 (ZGT4).

*Histological characterization and immunohistochemistry*

For confirmation of electrode placement animals were deeply anaesthetized with isoflurane 5% for induction and 1.5% for sustenance. Electrolytic lesions were made by passing an anodal current (30 µA for 10 s) through each tetrode, followed by a recovery period of 2 hours. Then animals were injected with 15 mg pentobarbital (i.p.) and transfused over the heart with 20 ml ice cold heparinized PBS followed by 30 ml 4% formalin. Brains were removed and post-fixed overnight in 4% formalin. They were then cryoprotected in 40% sucrose for 24–48 hours. Sections of 30 µm were cut in a cryostat and Nissl-staining with Cresyl violet was performed. A cover slip was placed on the slices with a mounting medium and then imaged on a confocal fluorescent microscope.

For quantification of PV cells, free-floating sections were washed in PBS plus 0.1% Triton X-100 (PBS-T) three times for 10 minutes each and then blocked by incubation with 10% Normal Donkey serum (NDS) in PBS-T for 1 hour. Free floating sections were incubated with primary antibodies for PV (Abcam 11 427; 1:500) for 24–48 hours at 4 °C in blocking solution containing 2% NDS. Sections were then washed in PBS-T, three times for 10 min each and then incubated with secondary antibody (Abcam: AB96947, 1:500) for 1 hour at room temperature. Slices were then mounted on glass slides and allowed to dry. A cover slip was placed on the slices with a mounting medium and then imaged on a confocal fluorescent microscope. Number of PV cells was counted using an 8bit image, automatically thresholded and analysed for number of particles using ImageJ software analysis tool.

*Determination of vigilance state*

We define wake episodes as periods of low amplitude EEG. Prominent theta band EEG activity and concurrent high amplitude EMG activity, corresponding to bursts of movement-related activity, and arousals shorter than 1 second were disregarded. Periods of low EMG tone with characteristic EEG and theta activity were scored as wakefulness including feeding and grooming behaviours. We defined NREM sleep as periods with a relatively high amplitude rich in low-frequency EEG and reduced muscle tone relative to wakefulness associated with behavioural quiescence. We scored REM sleep as sustained periods of theta band EEG activity and behavioural immobility associated with muscle atonia with brief phasic muscle twitches.

*Analysis*

Data analyses were carried out using custom scripts written in MATLAB® (R2018b, MathWorks, Natick, MA, USA). Furthermore, built-in functions from Wavelet and Signal Processing toolboxes of MATLAB were investigated.

*Spectral analysis*

Delta power in a recording segment was calculated using a modified periodogram with Hanning window (bandpower, MATLAB). Delta power during spindles was estimated using the periodogram with a 4-s Hanning window centered on spindles. Time–frequency representations were obtained using continuous wavelet transform with the complex Morlet function.

*Detection of spindles using an automated algorithm*

We detected spindles using the wavelet-based method proposed as previously published[31]. In brief, our criteria was using the wavelet energy time series which was smoothed using the 200 ms Hann window with a threshold equal to 3 SD (SD: standard deviation) above the mean and applied to detect potential spindle events. Events shorter than 400 ms or longer than 2 s were discarded. Using bandpass-filtered LFP signals in the spindle range (9–16 Hz), where we automatically counted the number of cycles of each event and excluded those with <5 cycles or more than 30 cycles. To discard artifacts we discarded those events with the power spindle band lower than 6–8.5 and higher than 16.5–20 Hz frequency bands (estimated spindle range). To discard false positives, the algorithm also measured the symmetry of spindles using the position of peak of wavelet energy time-series with regard to the start and end of spindles. False positives are discarded when the symmetry was lower than 0.5 and higher than 1 of the symmetric measure (between 0.5 and 1 values).

*Quantification of spindle rates during vigilance state transitions*

We estimated spindle rates before state switching from NREM to REM and wake, separately. We first marked all NREM–REM and NREM–Wake transition points, which were scored with 1 s resolution, and then calculated the spindle rates using different time scales, ranging from 5 to 40 s with a 5-s incremental window, before vigilance state transition. We averaged over all transitions for each animal to obtain the spindle rate (Figure 3 and Suppl. Figure 3).

*Correlation between slow waves and spindles*

We filtered LFP/EEG signals for SWs (0.5–4 Hz) and spindles (10–16 Hz) frequency bands using the 6000th and 333rd order window-based FIR filters, respectively, in both the forward and reverse directions. We extracted envelopes of spindles using the Hilbert transform. We then aligned both SWA and spindle envelope to the start of spindles, detected by algorithm, and averaged across entire NREM spindles. To quantify correlations between SWs and spindles, we estimated normalized cross-correlations between averaged signals of SWs and spindle envelopes. To find the ratio of spindles that coincide with UP states, we detected UP states as reported previously [19].

*Modulation Index*

We used the Modulation Index (MI) to measure phase–amplitude coupling [39]. We first bandpass-filtered LFP signals into low- and high-frequency bands, i.e. delta (0.5-4 Hz) and spindle (10-16 Hz), using finite impulse response (FIR) filters in both forward and reverse directions to eliminate phase distortion ("filtfilt" function, MATLAB). We designed FIR filters using the window-based approach ("fir1" function, MATLAB) with an order equal to three cycles of the low cutoff frequency. We then estimated instantaneous phase of low frequency and the envelope of high frequency oscillations using the Hilbert transform. Then, phase of the low frequency was discretized into 18 equal bins (N = 18, each 20°) and the average value of fast oscillations' envelope inside each bin was calculated. The resulting phase-amplitude histogram (P) was compared with a uniform distribution (U) using the Kullback–Leibler distance, DKL(P, U)=∑Nj=1P(j)∗log[P(j)/U(j)], which was normalized by log(N) to obtain MI, MI=DKL/log(N).

*Comodulogram analysis*

MI-based comodulogram analysis is a powerful tool to assess phase–amplitude coupling for a wide range of frequencies. The MI is independent of the power of fast frequencies and therefore the comodulogram graph is not biased for bands with a higher power. To explore coupling between different pairs of frequency bands, we considered 18 frequency bands for phase (0.5–20 Hz, 1-Hz increments, 2-Hz bandwidth), and 28 frequency bands for amplitude (20–310 Hz, 10-Hz increments, 20-Hz bandwidth). MI values were then calculated for all these pairs to obtain the comodulogram graphs. For each animal, we first filtered continuous 24-h recordings into the mentioned frequency bands and estimated the Hilbert transform. Then, we concatenated episodes of each vigilance state to derive the stage-specific comodulogram graph. To avoid power line interferences, frequency bands in the vicinity of 60 Hz and its harmonics were reassigned with a 2-Hz safe margin from these interfering frequencies.

*Time-resolved phase–amplitude coupling*

We used a time-resolved approach to track changes of phase–amplitude coupling in both time and frequency domains. We first bandpass-filtered the LFP/ signal into 24 sub-bands for amplitude frequency, as described in "Comodulogram Analysis" section, and obtained envelopes of filtered signals using the Hilbert transform. We segmented the resulting envelopes into 4-s windows having 75% overlap and calculated the fast Fourier transform (FFT) for each segment. To minimize the side effects of filtering on signal edges, we applied segmentation after filtering. The frequency band for the phase was obtained from the envelope of the filtered signal for the amplitude frequency band. We estimated the peak frequency of the envelope using the FFT, and then bandpass-filtered LFP signals at this peak frequency using a 2-Hz bandwidth. MI was then calculated between the phase of this frequency band and the corresponding high-frequency band. The logic behind this approach is that if there is phase–amplitude coupling between low and high-frequency bands, the modulating low frequency and dominant frequency of the envelope of the modulated fast frequency are similar. Therefore, we can track coupling dynamics over time and frequency by replacing phase-frequency axis in the comodulogram (horizontal axis) with the time axis.

*Single unit analysis*

Spike transitions were computed by averaging the spiking rate activity during the last 5 seconds before transition to another vigilant state and the averaged 5 seconds of spiking activity of the subsequent vigilant state per recorded site. Neurons that were not modulated by the sleep wake cycle were excluded in this analysis.

*Statistical methods*

MATLAB® (R2018b, MathWorks, Natick, MA, USA) and Prism 8 (GraphPad) were used for statistical analysis. No power calculations were performed to determine sample sizes, but similarly sized cohorts were used as in other relevant investigations [9,19]. Data were compared via two-way ANOVA followed by Bonferroni's multiple comparison test. Multiple comparisons tests for parametric data using comparisons between phenotypes (WT vs KO) and the different experimental conditions (baseline sleep vs recovery sleep), as indicated in the text. Values in the text are reported as mean ± standard error mean (SEM) unless reported otherwise. Figures were prepared in Adobe Illustrator CC (Adobe).

**Supplemental Figure legends.**

**Suppl. Figure 1**. Sleep characterization. A. Left: vigilante state amount, mean bout duration and Mean bout Wake ** P= 0.027, t=2.952, DF=21), Episode density (Wake dark WT-KO ***P <0.001, t= 6.20, DF=77.0); percentage vigilant state and mean bout duration: dark n=5,6; BL, 9,6; and SR: n=7,6 for WT and KO respectively. B-C. total percentage of wake and REM sleep during sleep recovery period (n=4 and 5 WT and KO respectively). Significant levels were calculated using 2- way ANOVA and Bonferroni's multiple comparisons test. All results are represented in mean +/- s.e.m. *P< 0.33, **P<0.002 and ***P< 0.001.

**Suppl. Figure 2.** Changes in in Slow wave activity and Delta oscillations on Gclm -/- KO mice. A. Representative micrograph of Cresyl violet staining showing the tetrode location. B. Normalized average data of Delta 1 (δ1) power (0.75-1.75) quantification during NREM sleep baseline (BL) and the first hour of sleep recovery time (SR) taken from the EEG and tetrodes in thalamocortical nuclei (EEG frontal (EEGfront) : P= 0.3749; t = 2.26; DF = 5.951, n=7 WT and n=5 KO; EEG parietal (EEGpar) : P= >0.9999; t = 0.2037; DF = 7.953, n=7 WT and n=5 KO; ACC : * P=0.0166, t= 5.327, DF 5.457, BL: n=7 WT and 5 KO; Brr : * P= 0.0135, t= 5.57, DF= 5.44, n=5 WT and n=4 KO; TRN : * P= 0.039; t=3.6, DF=9.093, n= 8 WT and n= 6 KO; VPL: ** P= 0.0018, t= 5.786, DF=9.09, n=7 WT and n=5 KO; AD : P= 0.8927; t = 1.177; DF = 8.021, n=7 WT and n=6 KO; SR n=7 WT and n=4 KO; Brr : * P= 0.0156, t=8.739, DF=3.271; VPL: ** P= 0.0089, t= 6.376, DF= 5.136). C. Positive slope of individual detected slow wave events during the light period in spontaneously occurring NREM sleep episodes (EEG front : **P= 0.004, t=3.63, n= 8,5; EEGpar : *P= 0.030, t=2.96,n= 8,5; ACC : ***P<0.001, t=4.15,n= 6,3; AD : n= 6,3; TRN : n=8,6; Brr : ***P < 0.001, t= 6.51, n= 8,6; and VPL : **P= 0.008, t= 3.41, n=7,6 for WT and KO respectively; DF= 62). During the dark, EEGfront : **P= 0.013, t=3.38, ,n= 4,5; EEGpar : *P= 0.041, t=2.94, n= 4,5; ACC: ***P<0.001, t= 4.41, n= 2,3; AD : *** P <0.001, t= 6.38, n= 2,2; TRN : n=4,3; Brr : *P= 0.043, t= 2.92, n= 4,3; and VPL : n= 4, 3 for WT and KO respectively; DF= 62). D. Delta 1 and 2 during the dark cycle. n values are the same as listed above. E. Positive slope of individual slow wave events, with n numbers as listed in C. All results are represented in mean +/- s.e.m. *P< 0.33, **P<0.002 and ***P< 0.001.

**Suppl. Figure 3**. Spindle activity during characterization. A. Normalized average sigma power (10-16 Hz) showing the individual values from before (black open circles) and after sleep deprivation during the first hour of sleep recovery (black filled circles) for WT and knock out (red). B. Spindle rate during calculated from the las 25 s of the NREM sleep till the transitions to REM sleep (EEGfront : n=8,5; EEG parietal (EEGpar) : n=8, 5; ACC :

n=6,3; AD : n=6,3; TRN : n=8, 6; Brr : n=8, 6; and VPL : n=7, 6 for WT and KO respectively). C. Representative detected (lines) spindle activity trough the sleep wake cycle during spontaneous occurring NREM sleep (light period), during the first hour of the sleep recovery (SR) and during the dark period during.

**Table1**. Spiking rate activity across sleep wake cycle. Data collected from spiking rate during all vigilant states: wake, NREM and REM sleep. Significant levels were calculated using 2-way ANOVA and Bonferroni's multiple comparisons test. All results are represented in mean +/- s.e.m. *P< 0.33, **P<0.002 and ***P< 0.001.

**Suppl. Figure 4.** Spiking rate during transition from different vigilant states. Summary data from all recorded sites during wake to NREM sleep (W2N), NREM to REM sleep (N2R), NREM sleep to wakefulness (N2W) and from REM sleep to wakefulness (R2W). ACC (W2N: ***P<0.001, t= 4.0; N2R *P= 0.016, t= 3.12, DF = 28, n = 5 and 5); AD (W2N: **P = 0.008, t = 3.69; N2R *P = 0.018, t = 3.27, R2W ***P < 0.001, t = 5.35, DF= 16, n = 2 and 4 for WT and KO); TRN (n = 15 and 13 for WT and KO); Brr (n =10, 7 for WT and KO); and VPL (n =10, 6 for WT and KO respectively). Significant levels were calculated using 2- way ANOVA and Bonferroni's multiple comparisons test. All results are represented in mean +/- s.e.m. *P< 0.33, **P<0.002 and ***P< 0.001.

*Suppl. Figure 1. Czekus C. et al*

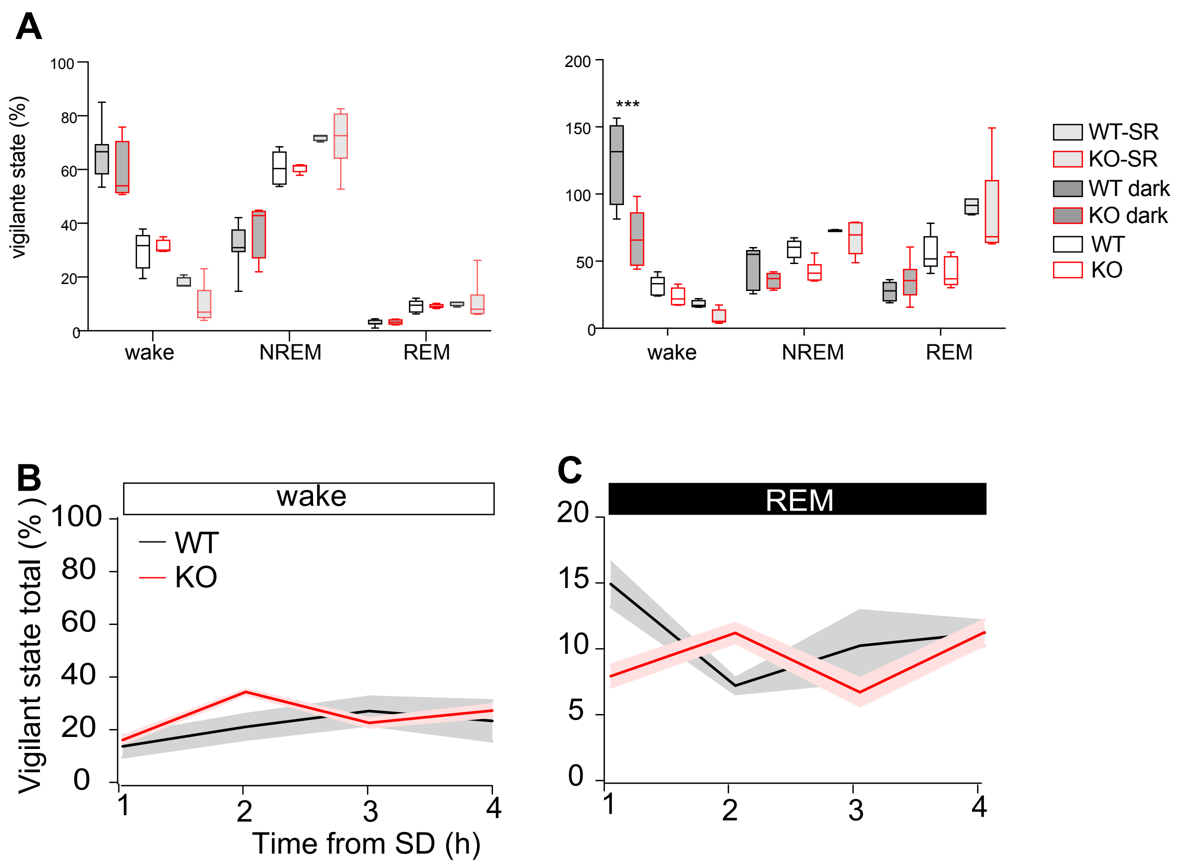

### Suppl. Figure 2. Czekus C. et al

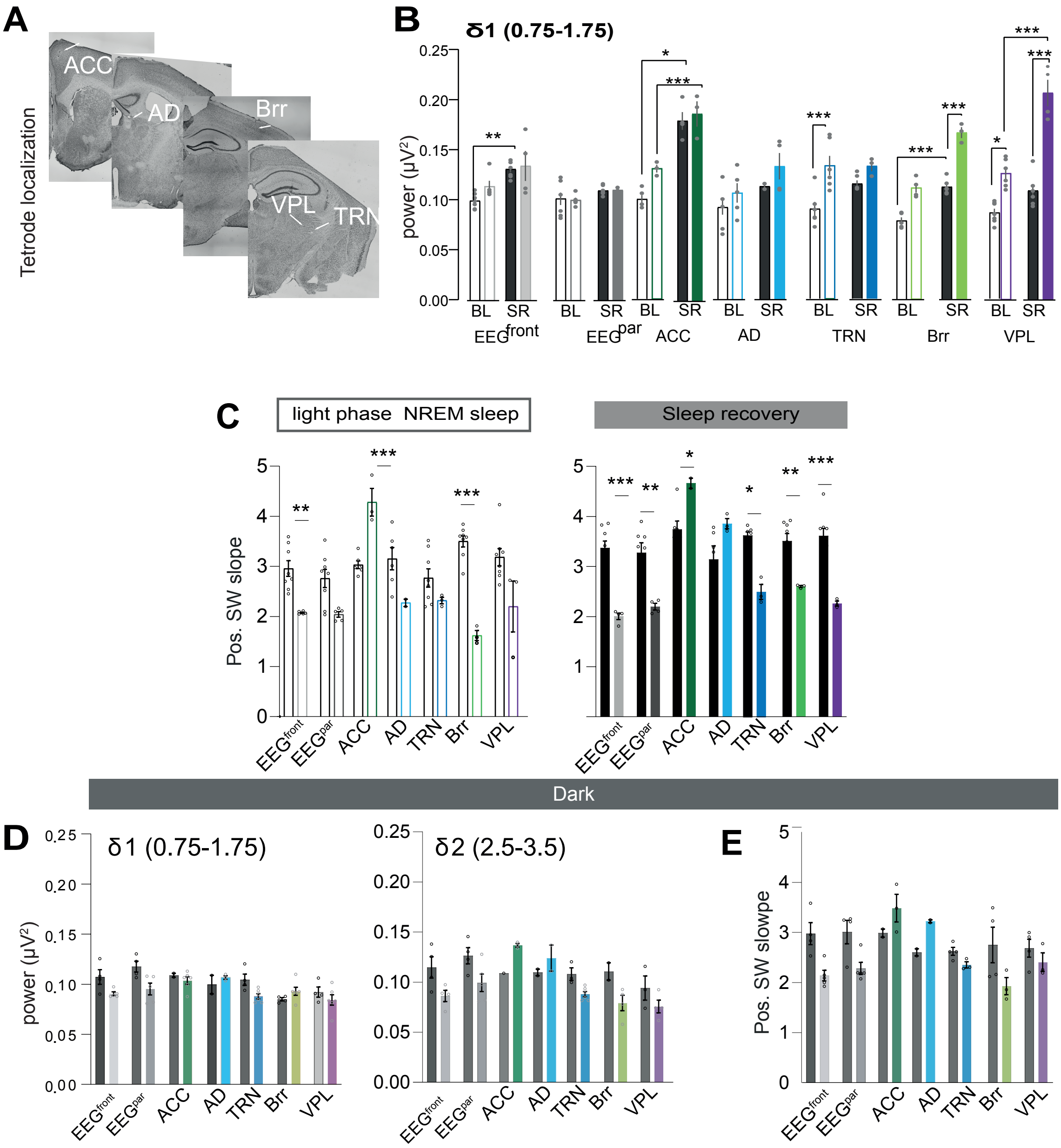

*Suppl. Figure 3. Czekus C. et al*

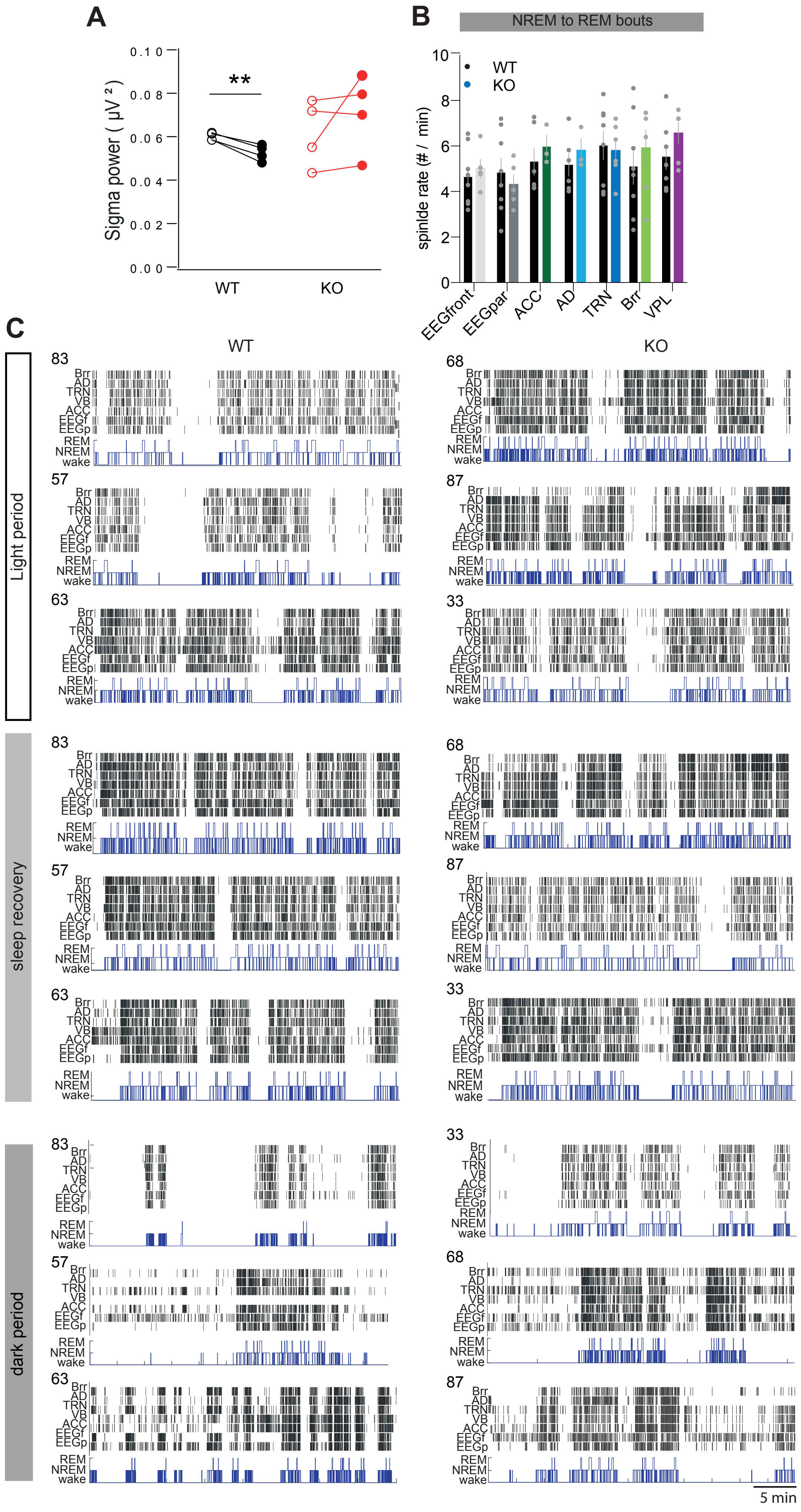

### Suppl. Table 1. Czekus C. et al

## A

| ACC | WT-BL | | | KO-BL | | |
|---|---|---|---|---|---|---|
| | Mean | SEM | N | Mean | SEM | N |
| wake | 23.04 | 4.34 | 5 | 31.25 | 5.58 | 6 |
| NREM | 5.37 | 0.49 | 5 | 13.24 | 1.95 | 6 |
| REM | 10.53 | 1.17 | 5 | 16.69 | 2.36 | 6 |
| | WT-SR | | | KO-SR | | |
| | Mean | SEM | N | Mean | SEM | N |
| wake | 21.83 | 8.59 | 4.00 | 31.82 | 0.23 | 3 |
| NREM | 8.52 | 0.77 | 4.00 | 10.03 | 0.37 | 3 |
| REM | 12.49 | 1.00 | 4.00 | 9.17 | 0.68 | 3 |
| Brr | WT-BL | | | KO-BL | | |
| | Mean | SEM | N | Mean | SEM | N |
| wake | 33.59 | 6.68 | 10 | 23.03 | 7.30 | 7 |
| NREM | 10.03 | 1.59 | 10 | 6.32 | 1.80 | 7 |
| REM | 19.05 | 3.20 | 10 | 11.48 | 4.05 | 7 |
| | WT-SR | | | KO-SR | | |
| | Mean | SEM | N | Mean | SEM | N |
| wake | 21.83 | 8.59 | 4 | 31.82 | 0.23 | 3 |
| NREM | 8.52 | 0.77 | 4 | 10.03 | 0.37 | 3 |
| REM | 12.49 | 1.00 | 4 | 9.17 | 0.68 | 3 |
| TRN | WT-BL | | | KO-BL | | |
| | Mean | SEM | N | Mean | SEM | N |
| wake | 42.00 | 6.06 | 18 | 23.04 | 4.53 | 14 |
| NREM | 22.31 | 3.60 | 18 | 8.46 | 1.53 | 14 |
| REM | 30.16 | 6.22 | 18 | 12.38 | 3.23 | 14 |
| | WT-SR | | | KO-SR | | |
| | Mean | SEM | N | Mean | SEM | N |
| wake | 27.70 | 6.36 | 8 | 42.96 | 6.70 | 9 |
| NREM | 14.50 | 2.96 | 8 | 9.43 | 0.62 | 9 |
| REM | 19.31 | 4.98 | 8 | 11.89 | 2.70 | 9 |
| VPL | WT-BL | | | KO-BL | | |
| | Mean | SEM | N | Mean | SEM | N |
| wake | 32.67 | 6.57 | 10 | 41.99 | 3.33 | 8 |
| NREM | 13.85 | 2.71 | 10 | 11.91 | 4.00 | 8 |
| REM | 17.31 | 4.12 | 10 | 24.59 | 5.96 | 8 |
| | WT-SR | | | KO-SR | | |
| | Mean | SEM | N | Mean | SEM | N |
| wake | 35.10 | 14.06 | 4 | 47.12 | 1.26 | 5 |
| NREM | 10.31 | 3.19 | 4 | 7.94 | 1.15 | 5 |
| REM | 26.58 | 10.91 | 4 | 16.58 | 3.27 | 5 |
| AD | WT-BL | | | KO-BL | | |
| | Mean | SEM | N | Mean | SEM | N |
| wake | 10.46 | 0.92 | 5 | 26.18 | 4.28 | 7 |
| NREM | 3.26 | 0.49 | 5 | 7.12 | 0.79 | 7 |
| REM | 6.29 | 0.99 | 5 | 14.03 | 2.94 | 7 |
| | WT-SR | | | KO-SR | | |
| | Mean | SEM | N | Mean | SEM | N |
| wake | 9.59 | 2.94 | 4 | 38.42 | 2.03 | 6 |
| NREM | 2.38 | 1.59 | 4 | 6.23 | 1.00 | 6 |
| REM | 3.33 | 1.97 | 4 | 19.79 | 3.38 | 6 |

## B

| WT - KO BL | | | | |
|---|---|---|---|---|
| | Summary | P Value | t | DF |
| ACC | * | 0.04 | 3.915 | 5.613 |
| Brr | ns | 0.24 | 1.611 | 14.84 |
| TRN | ** | 0.009 | 3.64 | 17.86 |
| VB | ns | 0.44 | 0.7987 | 17.32 |
| AD | ns | 0.08 | 3.361 | 4.195 |

| WT - KO BL | | | | |
|---|---|---|---|---|
| | Summary | P Value | t | DF |
| ACC | ns | >0.9999 | 0.434 | 47 |
| Brr | ns | >0.9999 | 0.6166 | 47 |
| TRN | ns | 0.1331 | 2.289 | 47 |
| VB | ns | >0.9999 | 0.7753 | 47 |
| AD | ns | 0.9857 | 1.308 | 47 |

| Wake BL | | | | |
|---|---|---|---|---|
| WT - KO BL | | | | |
| | Summary | P Value | t | DF |
| ACC | ns | >0.9999 | 0.758 | 75 |
| Brr | ns | >0.9999 | 1.197 | 75 |
| TRN | ** | 0.005 | 3.426 | 75 |
| VB | ns | >0.9999 | 1.098 | 75 |
| AD | ns | 0.6886 | 1.5 | 75 |

| Wake SR | | | | |
|---|---|---|---|---|
| WT - KO BL | | | | |
| | Summary | P Value | t | DF |
| ACC | ns | >0.9999 | 0.8198 | 47 |
| Brr | ns | >0.9999 | 1.174 | 47 |
| TRN | ns | 0.2744 | 1.969 | 47 |
| VB | ns | >0.9999 | 1.124 | 47 |
| AD | * | 0.037 | 2.8 | 47 |

| REM BL | | | | |
|---|---|---|---|---|
| WT - KO BL | | | | |
| | Summary | P Value | t | DF |
| ACC | ns | >0.9999 | 0.6526 | 80 |
| Brr | ns | >0.9999 | 0.9854 | 80 |
| TRN | ** | 0.0099 | 3.2 | 80 |
| VB | ns | >0.9999 | 0.9849 | 80 |
| AD | ns | >0.9999 | 0.8475 | 80 |

| REM SR | | | | |
|---|---|---|---|---|
| WT - KO BL | | | | |
| | Summary | P Value | t | DF |
| ACC | ns | >0.9999 | 0.3936 | 52 |
| Brr | ns | >0.9999 | 0.1579 | 52 |
| TRN | ns | 0.8651 | 1.382 | 52 |
| VB | ns | 0.81 | 1.419 | 52 |
| AD | ns | 0.1251 | 2.308 | 52 |

### Suppl. Figure 4. Czekus C. et al

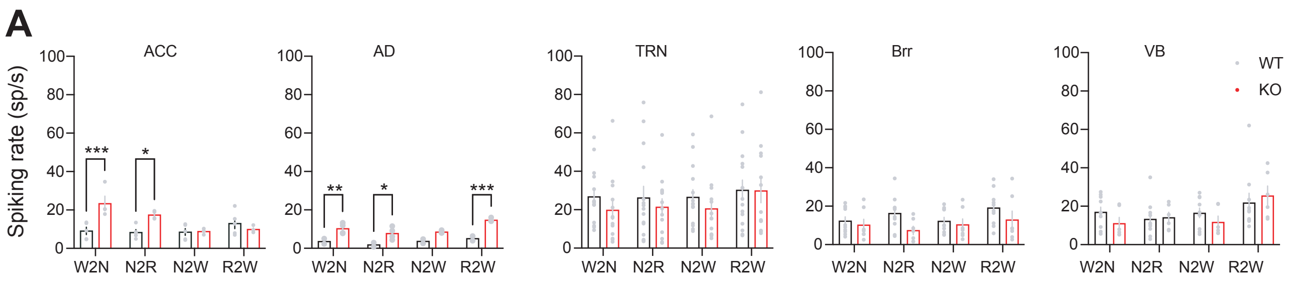
